# Bird occurrence and trophic interactions vary across gradients of tree diversity and microclimate in a planted forest

**DOI:** 10.1101/2025.02.17.637724

**Authors:** Sophie Coyne, Eriberto Osorio, Shelley K. Bennett, Aidan Kirchgraber, John D. Parker, A. Justin Nowakowski

**Author notes:** Open Research Statement: All data underlying analyses will be available in a publicly accessible repository upon acceptance. Author is currently affiliated with the Chesapeake Bay Trust under the Chesapeake Conservation and Climate Corps, 108 Severn Avenue, Annapolis, MD 21403.

## Abstract

Deforestation reshuffles communities across landscapes with myriad consequences for ecosystem function. Following deforestation, rapid exposure to novel microclimates can act as a strong environmental filter, favoring warm-adapted species and decoupling trophic interactions. Forest restoration may partly reverse this process through increased habitat structure, food resources, and buffering of microclimates – each potentially modified by tree diversity. Despite growing evidence that tree diversity and cool microclimates help maintain animal diversity in natural forests, less is known about how these factors shape species assemblages or multitrophic dynamics in restoration areas. Here, using surveys and two field experiments within a long-term tree planting experiment, we assessed the relative effects of tree diversity, forest structure, and associated microclimate on fine-scale space use by birds and their top-down impacts on insects. Surveys showed that fine-scale occurrences of birds increased in cooler plots, which were associated with higher tree diversity and structural complexity. The strength of microclimate effects on bird occurrences was strongest for species that are forest specialists. To assess risk to insect herbivores from avian predation, we used a sentinel prey experiment and found that predation risk increased in warmer plots, counter to our expectations based on bird surveys. Last, we examined top-town effects of bird exclusion on leaf herbivory, finding that skeletonizing patterns of herbivory increased in exclosures and in cooler plots. Taken together, these results suggest that microclimate resulting from variation in forest structure shapes space use of birds at fine scales with complex outcomes for bird-herbivore-tree interactions in planted forests. Active restoration methods that enhance below-canopy cooling may improve biodiversity outcomes and help maintain species interactions that underlie many ecosystem functions.

## INTRODUCTION

Despite covering about 30% of the Earth’s surface, forests support most of the terrestrial biodiversity on the planet (FAO and UNEP 2020; Pillay et al. 2022). However, human population growth and associated land use changes have led to widespread deforestation and forest fragmentation worldwide (FAO and UNEP 2020). The resulting habitat loss is driving the global biodiversity crisis, which is further exacerbated by increased intensity and frequency of extreme temperatures under global climate change (IPBES 2019; Jaureguiberry et al. 2022; Newbold et al. 2015). Reforestation has become a leading intervention for increasing resilience to climate change, restoring ecosystem function, and increasing biodiversity in deforested areas (e.g., Target 2 of the Kunming-Montreal Global Biodiversity Framework 2022). Forest restoration can increase local biodiversity by 15-84% on average (Aerts and Honnay 2011; Crouzeilles et al. 2016). However, determining the role of diversity and composition of planted trees as well as the biotic and abiotic mechanisms leading to successful forest restoration remain active areas of research (Aerts and Honnay 2011; De Frenne et al. 2021; IPBES 2019).

The diversity of trees in restoration areas may be important for re-establishing ecosystem dynamics by facilitating the recovery of animal diversity and associated interactions in reforestation projects (Depauw et al. 2024). Natural forests with greater levels of tree species diversity better support ecosystem processes through community assembly dynamics (Vehviläinen, Koricheva, and Ruohomäki 2007; Ampoorter et al. 2020; Wildermuth et al. 2023) and resulting multi-trophic interactions (Alavez et al. 2023; Vázquez-González et al. 2024).

Increased tree species diversity is generally associated with increases in animal species diversity, as shown in previous studies of bird and arthropod assemblages (Barbaro et al. 2019; Ampoorter et al. 2020; Butz et al. 2023; Vázquez-González et al. 2024). Other studies show that increased functional diversity of trees can mediate the impacts of insect herbivores on trees, an important trophic relationship that affects tree growth and survival (Jactel and Brockerhoff 2007; Espelta et al. 2020; Wildermuth et al. 2023). Less is known about responses of multiple trophic levels to tree diversity in forest restoration projects or the pathways that underlie the effect of tree diversity on animal diversity (Depauw et al. 2024).

Recent work shows that microclimates can play a central role in shaping animal distributions across landscapes and within forests, filtering assemblages and likely altering species interactions and associated ecosystem functions (Frey, Hadley, and Betts 2016; Nowakowski, Frishkoff, et al. 2018; Williamson et al. 2022). In deforested areas, daytime air temperatures near the ground can be >10 C greater than nearby forests, exceeding the physiological tolerances of some species but not others (Nowakowski, Frishkoff, et al. 2018). As forests grow in age and complexity, they are able to sustain stable microclimates that are on average cooler than in deforested areas during the day and in the summer, and warmer at night and in winter, as a result of changes in the vertical and horizontal structure of forest canopies and understories (Chen et al. 1999; Potter, Teclaw, and Zasada 2001; von Arx et al. 2013; Z. Zhang et al. 2021). These cooler microclimates buffer many species from extreme temperatures and can facilitate local persistence (Frey, Hadley, and Betts 2016; Nowakowski, Watling, et al. 2018; De Frenne et al. 2021).

For climate-sensitive insect predators, such as many forest birds, microclimate refugia may help stabilize long term occupancy dynamics and breeding activity in spite of ongoing shifts in macroclimate suitability (Frey, Hadley, and Betts 2016; Kim et al. 2022). The impacts of microclimate can be a stronger driver of bird population dynamics than vegetation structure and composition but vary among bird species (Frey, Hadley, and Betts 2016), possibly as a function of their thermal physiology and macroclimate niches (Frishkoff et al. 2016; Nowakowski, Frishkoff, et al. 2018). For ectotherms, high temperatures and low humidity within deforested areas lead to increased metabolic rates and water loss, extirpating some species as they are exposed to conditions near their physiological limits (Lemoine, Burkepile, and Parker 2014; Nowakowski, Watling, et al. 2018; Wilson and Fox 2021; Williamson et al. 2022). Within arthropod communities, microclimate gradients can act as a strong community filter, with cooler microclimates often supporting greater community diversity and abundance (in temperate and tropical, low-mid elevation regions (Williamson et al. 2022; Aggarwal et al. 2023)). Despite growing evidence of multi-taxa reliance on cooler microclimates in forests, there is little research on how microclimate may impact species assemblages or multitrophic dynamics in restoration areas.

In particular, animal diversity and multi-trophic interactions can serve as important outcomes for evaluating forest restoration success, and understanding the impact of microclimate on these processes may aid in forest restoration design. For example, bird-insect-tree interactions can shape changes in forest composition and structure over time. Insectivorous birds impact insect populations in forests, resulting in top-down effects on herbivory across many tree species and forest types (Holmes, Schultz, and Nothnagle 1979; Marquis and Whelan 1994; Barber and Marquis 2009; Bridgeland et al. 2010; Mäntylä, Klemola, and Laaksonen 2011; Forde et al. 2022; Getman-Pickering et al. 2023; 2023). Greater insect herbivory can reduce growth and survival of saplings and seedlings (Giffard et al. 2012) and affect biomass of mature trees (Marquis and Whelan 1994; Boege and Marquis 2006; Bridgeland et al. 2010; Mäntylä, Klemola, and Laaksonen 2011). Predation of herbivorous insects by birds can generate positive top-down effects on tree biomass, reduce mortality rates, and reduce leaf damage (Marquis and Whelan 1994; Mooney et al. 2010; Mäntylä, Klemola, and Laaksonen 2011; Getman-Pickering et al. 2023). If tree diversity and/or buffered microclimates in planted forests promote more diverse and abundant assemblages of insectivorous birds, these dynamics could lead to reduced herbivory and in turn influence changes in forest structure and composition over time.

In this study, we address the overarching question of how tree diversity, forest structure, and associated microclimate each affect variation in trophic interactions at fine scales in a forest restoration experiment. To do so, we first examined the frequency of occurrence of bird species across gradients of diversity, forest structure, and associated microclimates in plots planted with different tree diversities and species compositions. Occurrence within plots represents fine-scale patterns of space use by birds, which may depend on their landscape-scale habitat affinities (i.e., forest specialist, generalists, and field specialists). We then asked whether predation risk for herbivorous insects by birds also varies across diversity, structure, and microclimate gradients. Finally, we examined whether the extent of leaf damage from herbivory and top-down effects of bird predation on herbivory varied across these gradients. We addressed these questions by collecting observational data on bird occurrence, foraging behavior, and microclimate in a tree diversity restoration experiment in Maryland, USA. We also conducted a bird-exclusion experiment to measure top-down effects on herbivory. We hypothesized that greater diversity in vegetation and cooler microclimates would be associated with increased bird abundance and species richness at fine spatial scales. While birds may be both cool or warm adapted, we predicted that bird occurrence and foraging activity would generally be positively associated with cooler microclimates during the summer. From this, we expected that insect predation by birds would increase within cooler microclimates and more diverse plots, leading to greater top-down effects on herbivory.

## METHODS

### Study Area

We conducted this study within the BiodiversiTREE experiment at the Smithsonian Environmental Research Center (SERC) in Edgewater, Maryland, USA (38°52’ N, 76°33’ W). BiodiversiTREE is a tree diversity experiment that was established on 24 ha of former cropland in April 2013 using a biodiversity ecosystem function (BEF) design to investigate forest response to restoration and climate change. The 16 focal species in BiodiversiTREE were selected from some of the most common species by basal area in the focal region and include *Acer rubrum*, *Carpinus caroliniana*, *Carya alba*, *C. glabra*, *Cornus florida*, *Fagus grandifolia*, *Fraxinus pennsylvica*, *Liriodendron tulipifera*, *Liquidambar styraciflua*, *Nyssa sylvatica*, *Platanus occidentalis*, *Quercus alba*, *Q. pagoda*, *Q. rubra*, *Q. velutina*, and *Ulmus americana*. Trees were planted in 35m x 35m plots as either species monoculture plots (*n* = 32, 2 plots per species), 4-species plots (*n* = 19), 12-species plots (*n* = 19), or unplanted natural regeneration plots (*n* = 5), totaling 75 plots (see (Griffin et al. 2019; Devaney et al. 2020) for more information on site design). Differences in tree diversity and composition among plots generate substantial variation in tree height, canopy cover, and forest structure that in turn contribute to microclimate gradients. The BiodiversiTREE site has a warm-temperate climate and was previously planted (>35 years) with corn (*Zea mays* L).

### Bird point counts

To characterize patterns of bird occurrence across gradients of tree diversity, forest structure, and microclimate, we conducted standardized point counts in a subset of 49 BiodiversiTREE plots, including monoculture plots (n =16), 4-species plots (n=17), and 12-species plots (n=16). We conducted three rounds of point counts in all study plots, each over approximately eight days, from June 22 - 30 2023, July 10 - 18 2023, and June 26 - August 2, 2024. We surveyed a subset of plots that spanned levels of tree diversity to ensure that each round of point counts could be completed in a short period (approx. a week or less). All point counts were completed after peak migrations in this region, which usually start in mid to late May (Hagan, Lloyd-Evans, and Atwood 1991). For each day of sampling, the first plot surveyed was randomly selected, and adjacent plots were never sampled sequentially to reduce potential disturbance caused by the observer. Counts were conducted between 0600 and 0900 hrs when many birds are most active (Verner and Ritter 1986; Ellis and Taylor 2018). An observer stood in the center of the plot and recorded bird activity and sightings within the individual plot for six-minute periods, with a two-minute waiting period before starting each point count to reduce potential of including interruptions of bird activity into surveys (Thibault and Elmendorf 2024). Due to the small size of plots, observed birds were counted only by visual sighting or when audibly heard directly above to account for species that prefer the higher canopy. Distance of the bird from the observer was recorded with a laser range finder. Bird activity such as singing, calling, flying, or foraging was also recorded.

To examine whether landscape-scale macrohabitat affiliations predict fine-scale space use, we used bird point count data from NEON to classify birds observed in BiodiversiTREE plots according to their macrohabitat affiliations at the landscape scale (Thibault and Elmendorf 2024). We compiled point count data from 24 NEON plots across the SERC landscape (1072 ha) sampled each year from 2018-2022. We then estimated the difference in abundances of each species in forests and fields (see methods below) and classified species into three habitat affiliation categories. These three classifications included field specialists for species more likely to be found in field-like environments like croplands, forest specialists more likely to be found in forested environments, and generalists that were equally likely to be found in both fields and forests.

### Measuring forest structure and microclimate

We characterized tree diversity, stem density, canopy structure, topography (slope), and near-ground microclimate temperatures within each plot. We included multiple variables in our analyses that capture complementary dimensions of forest structure - canopy height, vertical complexity, and stem density - along with slope because vegetation structure and topography are known to be important microclimate forcing factors and can also influence animal assemblages. Tree diversity and density were based on plot treatments and vegetation surveys, respectively.

Although tree density was standardized at the time of planting, density varied across time due to variation in survival that was not associated with tree diversity (King et al. 2023). We characterized structural and topographic metrics using LiDAR data from 2022 collected by the NEON Airborne Observation Platform. We calculated slope in degrees from the level 3 digital terrain model product at 1 m resolution. Using LiDAR point clouds, we calculated a suite of structural diversity metrics using the lidR package in R that characterized canopy structure and vertical complexity within plots. We first filtered outlier and duplicate points and normalized points by ground height using spatial interpolation based on a Delaunay triangulation. Many structural metrics from LiDAR are highly correlated. Therefore, we chose two metrics that we expected to be important dimensions for characterizing bird habitat - canopy height and vertical complexity index; the latter characterizes the distribution of points across vertical bands above 0.5 m. We measured air temperatures every 15 min at 15 cm above the ground using TOMST TMS-4 sensors placed in the center of each plot. To characterize spatial variation in near-ground microclimate conditions across plots in the summer, we calculated the mean and 95th percentile of daytime temperatures in each plot for the period of June 1, 2023 to July 18, 2023.

### Foraging Experiment

We assessed predation risk for herbivorous insects across plots by conducting a sentinel prey experiment (Lövei and Ferrante 2017; Kolkert et al. 2021). A total of 36 feeding stations were constructed using six-inch diameter circular planter saucer dishes fastened atop 1.2 m tall wooden fence posts. We installed two feeding stations each within 18 plots (6 per diversity treatment) that were a subset of 49 plots used in the bird point counts. A Bushnell Core DS No Glow Trail Camera was aimed at each feeding station and motion triggered to capture photos and videos during day and night. Camera traps were set to take two photos and record a 10 second video directly after detecting movement near feeding stations. As predation rates are typically higher with live sentinel prey than artificial prey like plasticine models (Lövei and Ferrante 2017; Zvereva and Kozlov 2023), we used mealworms (*Tenebrio molitor*) as sentinel prey in feeding stations. Fifteen mealworms were placed on each dish and were replenished daily from July 20 - August 2, 2023. Each day, we counted the number of missing mealworms at each feeding station. We recorded the species and time of interaction for any birds and mammals that were captured by cameras interacting with the feeding station or eating mealworms.

### Bird exclosure experiment

To examine the potential top-down influence of bird predation on rates of insect herbivory across BiodiversiTREE plots, we conducted an exclosure experiment using a before, after, control, impact (BACI) design. For this experiment, we selected three focal tree species. For each species we selected six plots (2 plots per tree diversity treatment), with 18 total plots. Within each plot, we selected two individual trees of the focal species in the outer two rows of each plot that were a minimum of 5 m apart for inclusion in the exclosure experiment. While multiple focal species may have been present in each polyculture plot, only one species was chosen for study in each of the selected plots. The focal species were white oak (*Quercus alba*), sweetgum (*Liquidambar styraciflua*), and American beech (*Fagus grandifolia*). We selected both early (*L. styraciflua*) and later successional (*Q. alba and F. grandifolia*) species for experimentation. Successional status was considered as this greatly impacts the level of tree growth and canopy cover within plots, thus modulating microclimate. For example, *F. grandifolia* and *Q. alba* (late successional) monoculture plots were characterized by short stature trees and low canopy cover, compared to *L. styraciflua* monocultures that had larger trees and mostly closed canopies.

On each tree, we selected one branch as the treatment branch (exclosure applied) and a second, similarly positioned and sized branch was selected as a control branch (no exclosure) (*n = 72* branches total). We built exclosures to cover specific tree branches to exclude insectivorous birds from consuming insect herbivores (Holmes, Schultz, and Nothnagle 1979; Hooks, Pandey, and Johnson 2003; Boege and Marquis 2006; Garibaldi et al. 2010; Mäntylä, Klemola, and Laaksonen 2011; Giffard et al. 2012). For the body of the exclosures, we used green, 35.5 x 35.5 x 83 cm rectangular tomato trellises with 25.4mm x 25.4mm green, nylon mesh wrapped around the entire structure (Maas et al. 2019). The netting was wide enough to allow free transit of insects and arachnids while also being small enough to prevent access by the smallest insectivorous birds noted in the plots, including Carolina Wren (*Troglodytes ludovicianus*) and Blue-gray Gnatcatcher (*Polioptila caerulea*). Exclosures were left on the trees for four weeks (July 5 – July 31, 2023).

We estimated percent herbivory on selected leaves of study branches just prior to application and immediately following the removal of the exclosures. Each observer analyzed the same branches pre- and post-exclosure application. However, to control for potential observer bias, the post-exclosure estimation of herbivory was completed blind, such that observers were not aware of whether a branch was the treatment or control branch. Prior to application of exclosures, we quantified the total number of leaves on each branch (study and control) and 30 were selected for study. If a branch had < 60 leaves, the first 30 leaves from the end of the branch were sampled. If a branch had ≥ 60 leaves, then every *nth* leaf from the end moving up the branch was selected until 30 leaves were chosen (adapted from Boege & Marquis, 2006). For example, if a branch had 90 leaves, every third leaf would be chosen. Visual inspection followed the HerbVar Damage Estimation Guidelines (Ian Pearse et al. 2021).

Damage was visually estimated and recorded by type, here focusing on skeletonizing and chewing damage. We recorded damage for each category as percentages to the nearest 2.5% if covering more than 10% of leaf area, to 1% for levels between 1% and10%, and to 0.1% for levels between 0.1% and 0.9% (however, these values were aggregated prior to analysis - see methods below). We also calculated total insect damage per leaf, inclusive of skeletonizing and chewing patterns of herbivory, because they reflect damage by larger insects (e.g., caterpillars) that represent important prey or many bird species.

### Statistical Analyses

To assess the drivers of bird occurrence at fine spatial scales among BiodiversiTREE plots, we used piecewise structural equation modeling (SEM; (Lefcheck 2016)). We estimated standardized coefficients to quantify the relative importance of pathways among environmental variables measured for each plot, including tree diversity, forest structural metrics, slope, and microclimate, and their contributions to variation in bird occurrences. We focused on two response variables, the occurrences of any bird within the plot and the number of species observed during point counts. Therefore, component models of the piecewise SEM were generalized linear models where bird occurrence or number of observed species was the response, fit with binomial and Poisson distributions, respectively, and linear models where environmental variables were the response. Prior to fitting models, we assessed correlations among predictors and variable inflation factors, which were <5 for all variables.

To assess whether macrohabitat affiliations affect fine-scale space use, we first classified birds observed in BiodiversiTREE plots according to their macrohabitat affiliations at the landscape scale using NEON bird point count data. To do so, we fit a Bayesian generalized linear mixed model (GLMM), specifying a Poisson probability distribution, to estimate the mean difference in the number of observations per species per NEON plot between sites in fields versus forest. We fit a model with an intercept only for the main effects and a varying slope term among species for the effect of habitat type. These varying slope estimates represented the log factor change in abundance from field to forest habitat for each species. We then classified species as field specialists and forest specialists using the factor abundance change threshold of 0.5 and 1.5x, respectively, based on model estimates for each species. Species that were >1.5x more abundant in forest than field habitat were considered forest specialists. Species that were <0.5x less abundant in forest than in field habitat were classified as field specialists, and species that fell between those thresholds were classified as generalists. Models were fit in Stan using the brms package to interface with the R environment. We specified normal, uninformative priors.

We ran two chains with a total of 5000 iterations, sampled every 20 iterations, and with 500 iterations discarded as burnin. We ensured the chains had converged and mixed based on trace plots and the Gelman-Rubin statistic (all ∼1.0). Finally, we used these classifications to examine the effect of the interaction between microclimate and habitat affiliation on probability of bird occurrence and observed numbers of species in BiodiversiTREE plots using GLMs.

We then analyzed the rates that sentinel prey were depleted from feeding stations to assess predation risk by birds across tree diversity, structure (canopy height and vertical structure), and microclimate gradients in a subset of 18 BiodiversiTREE plots. Because heavy rains washed mealworms from feeding stations during several days of the experiment, we only analyzed counts of missing mealworms following days with no rain. We assessed relative importance of each variable by comparing effect sizes from a Bayesian GLMM fit with all explanatory variables. During the experiment, it was apparent that the number of missing mealworms and number of feeding stations with missing mealworms increased over time, likely as predators became habituated to the feeding stations. Therefore, we included the interaction between the day of the experiment and each environmental variable in the model. These environmental variables included the 95th percentile of daytime air temperatures, canopy height, vertical complexity, and tree diversity in plots. We specified a Poisson probability distribution to model counts of missing mealworms each day of the experiment and accounted for repeated measures within plots by including plot as a varying intercept term. Number of iterations and priors were defined as above. We examined photos and videos from July 22, July 24, and July 27 to summarize which species interacted with the feeding stations.

To analyze the effect of bird exclusion on herbivory, we used Bayesian implementation of a GLMM, specifying a Bernoulli probability distribution to analyze binary herbivory outcomes (herbivory or no herbivory per surveyed leaf). Although we estimated percent herbivory per leaf, nearly half of the observations were 0s (no herbivory), and, for non-zero values, most percent herbivory observations were <= 8% (median = 2%) - at the margin of precision for visual estimation. Therefore, we focus our analyses and inferences on probability of leaf herbivory modeled from binary outcomes. For each model, we included the specific tree branch being analyzed as a random intercept term to control for non-independence of multiple leaves counted on the same branch and repeated measurements of branches (before and after exclosure placement). As the response variable, we analyzed total observed insect herbivory, combining herbivory attributable to chewing and skeletonizing patterns of herbivory. We also analyzed chewing and skeletonizing types of herbivory separately. We combined the variables of control/treatment and before/after exclosure application into one composite BACI variable with four factor levels to test for treatment effects before and after placement of exclosures.

We then fit models to quantify the overall effects of exclosure treatment, tree species, tree diversity, canopy height, microclimate, and predation risk on leaf herbivory. Predation risk was based on the number of bird detections at foraging stations from the sentinel prey experiment.

For each response variable, we first fit additive models that included all explanatory variables as additive terms, and we assessed the relative importance of these variables in explaining variation in herbivory based on statistical support and their effect sizes. Next, to avoid overfitting, we fit separate models with interaction terms that allowed us to assess the dependence of treatment effects on the environmental covariates listed above. Specifically, we fit one model with just the interaction between treatment and tree species, as field observations and data exploration showed that the amount of total herbivory depended heavily on tree species. We fit four additional models, each with the interaction between treatment and tree species as well as the interaction between treatment and one of the remaining environmental covariates (tree diversity, canopy height, air temperature, or bird detections at feeding stations). We then compared all models using leave-one-out information criterion (LOOIC) and report treatment effects from models with the lowest LOOIC values. All models were fit in STAN using the brms R package, with the same priors and numbers of iterations described above. Before conducting all sets of analyses, we assessed potential for multicollinearity by examining pairwise correlations among explanatory variables and variable inflation factors. All analyses were conducted in R version 4.2.3.

## RESULTS

### Effect of tree diversity and microclimate on fine-scale bird occurrence

After controlling for slope of terrain, forest structure, and tree diversity, microclimate had the strongest direct effect on fine-scale variation in bird occurrences across plots (Fig 1).

**Figure 1.**
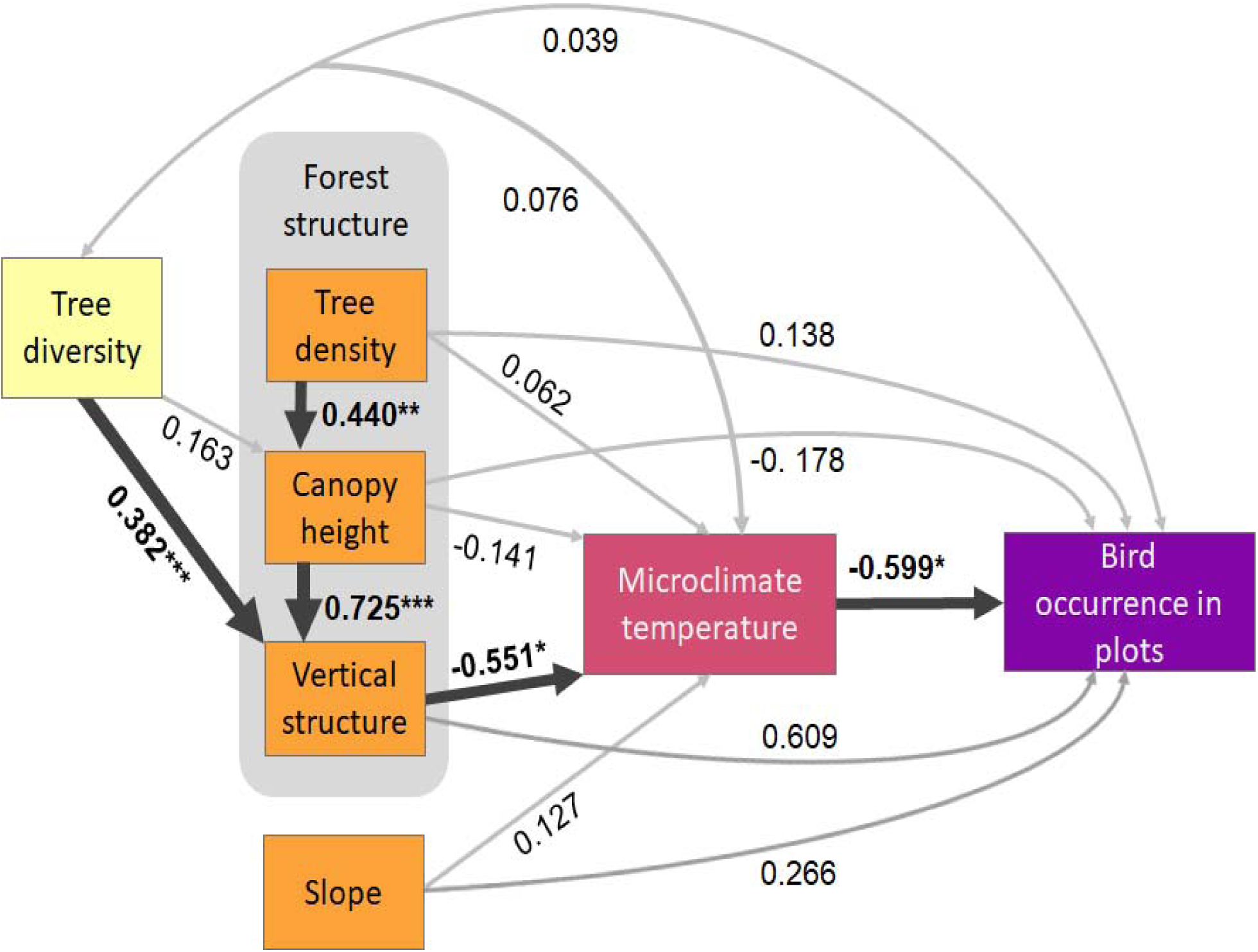
Microclimate was the primary driver of variation in bird occurrences across plots. Diagram shows hypothesized causal pathways among tree diversity, forest structure, slope, microclimate and probability of bird occurrences in forest plots. Values shown are standardized coefficients from a piecewise structural equation model. Black arrows indicate significant model estimates for a given path (\**P* < 0.05, \*\**P* < 0.01, \*\*\**P* < 0.001).

Specifically, the probability of bird occurrence significantly increased with decreasing daytime air temperatures within plots. Combined, all variables explained 24% of the spatial variation in bird occurrences. Greater vertical structure within plots was a significant driver of cooler microclimate temperature (*R*^2^ *=* 0.41 for combined predictors of microclimate), and greater tree diversity and height were associated with greater structural complexity of forest plots (*R*^2^ *=* 0.75 for predictors of vertical structure). Therefore, tree diversity indirectly influenced bird occurrences by enhancing forest structure and, consequently, cooling below canopy air temperatures (Goodness of fit test indicates key pathways have been included in the model, given the variable set: Chi-Squared = 3.719; *P* = 0.293). Model results were qualitatively similar when the number of observed bird species was included as the response variable, with the difference of marginally non-significant effects of temperature having the strongest effects on the number of species (Fig. 1; Appendix S1: Fig. S1).

We classified bird species observed in BiodiversiTREE plots according to macrohabitat affiliations based on their relative abundances in forest and field habitats across the landscape (using NEON data), which resulted in classification of four forest specialists, nine generalists, and five field specialist species (Fig. 2a). When grouping species by these habitat affiliations, there was a significant interaction indicating that the magnitude of the effect of microclimate on the probability of occurrence and number of species within plots depended on macrohabitat affiliations of species at the landscape scale (Fig. 2b and Appendix S1: Fig. S2; 95% Bayesian Credible Intervals exclude 0 for all interactions terms; Appendix S1: Table S1). Forest specialists exhibited the strongest, negative response to increasing air temperature while generalists exhibited an intermediate response, and field specialists had the weakest association with temperature.

**Figure 2.**
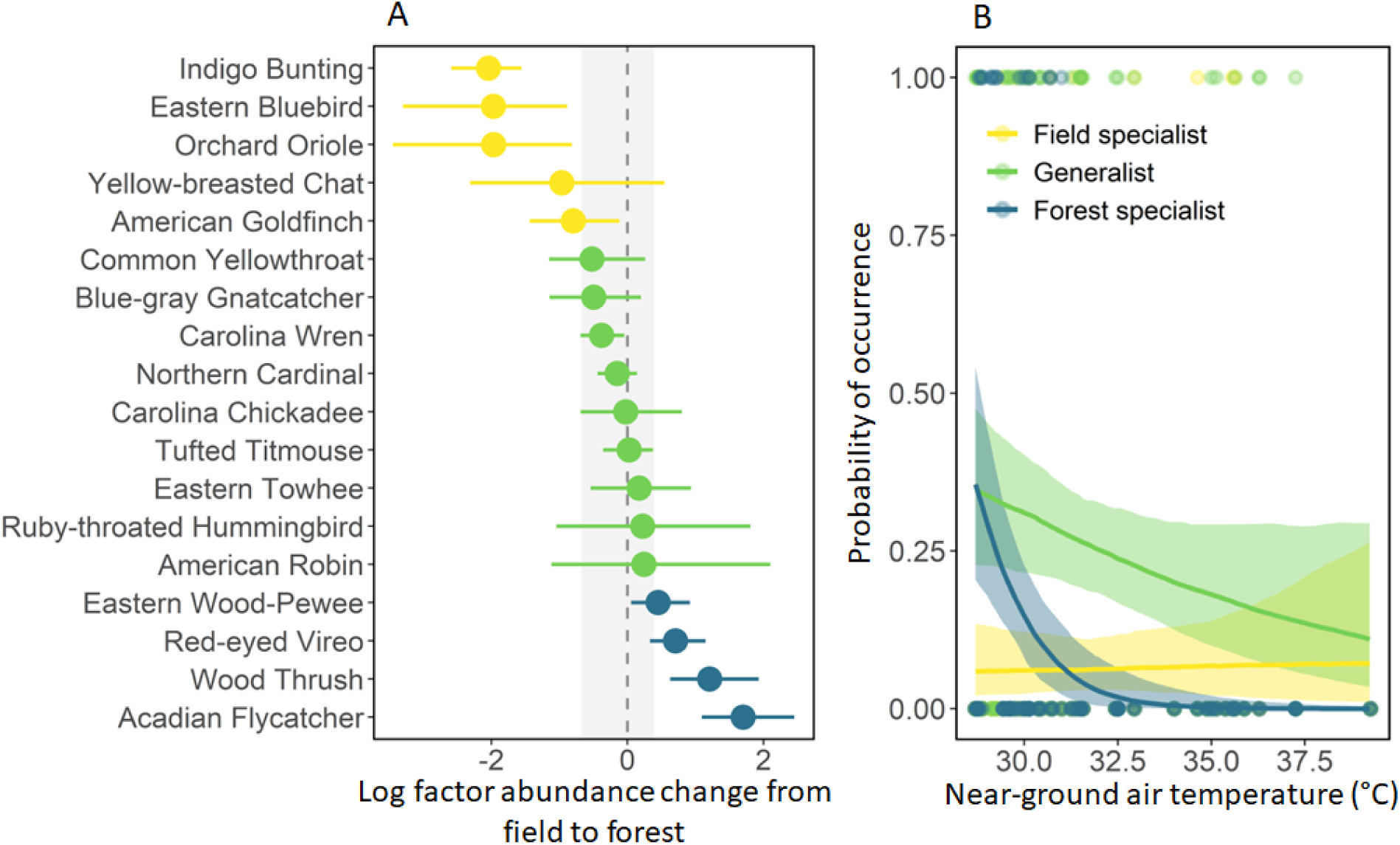
Responses of birds to microclimate depended partially on their macrohabitat affiliations. (A) Classification of bird species observed in point counts as field specialists (yellow), generalist (green), and forest specialists (blue) based on their relative abundances in NEON plots across the study landscape. Classifications are based on estimated differences in abundance from field to forest habitats from a generalized linear mixed model. The vertical grey band shows the thresholds used for classification - the natural log of a 0.5 and 1.5 factor abundance change for distinguishing field specialists and forest specialists, respectively, from generalist species. Error bars are 95% credible intervals. (B) The predicted effect of the interaction between landscape habitat affiliations and responses to microclimate variation at fine spatial scales across plots, from a generalized linear model. Bands indicate 95% credible intervals for predictions.

### Foraging behavior across forest structure and microclimate gradients

Rates of mealworm predation during the sentinel prey experiment increased significantly both over the duration of the experiment (β = 0.62, 95% Bayesian Credible Interval = [0.53, 0.73]) and across space, with increasing temperature (Appendix S1: Fig. S3; β = 0.27, 95% BCI = [0.04, 0.47]). There was a significant interaction between day of the experiment and air temperature (β = −0.09, 95% BCI = [−0.16, -0.03]; Appendix S1: Table 2), such that the effect of microclimate was greatest initially and then weakened over time, perhaps as predators became habituated to the feeding stations. No other explanatory variables were significant in the model (BCIs overlapped 0). Seven bird species were detected at mealworm stations, and frequency of detections in camera trap photos was greatest for field-associated and generalist species, especially Eastern Bluebirds (*Sialia sialis*), Tufted Titmouse (*Baeolophus bicolor*), and Carolina Wren (*Thryothorus ludovicianus*) (Fig. 3). There were 4 detections of raccoons (*Procyon lotor*) at feeding stations; therefore, the majority of detections at stations were of birds, and we infer that rates of mealworm depletion are largely attributable to these species. Despite the greater number of generalists and forest specialists detected in plots during point counts and the association of these species with cooler microclimates, bird foraging activity at stations was consistently higher in warmer plots and attributable, disproportionately, to field specialists and generalist species.

**Figure 3.**
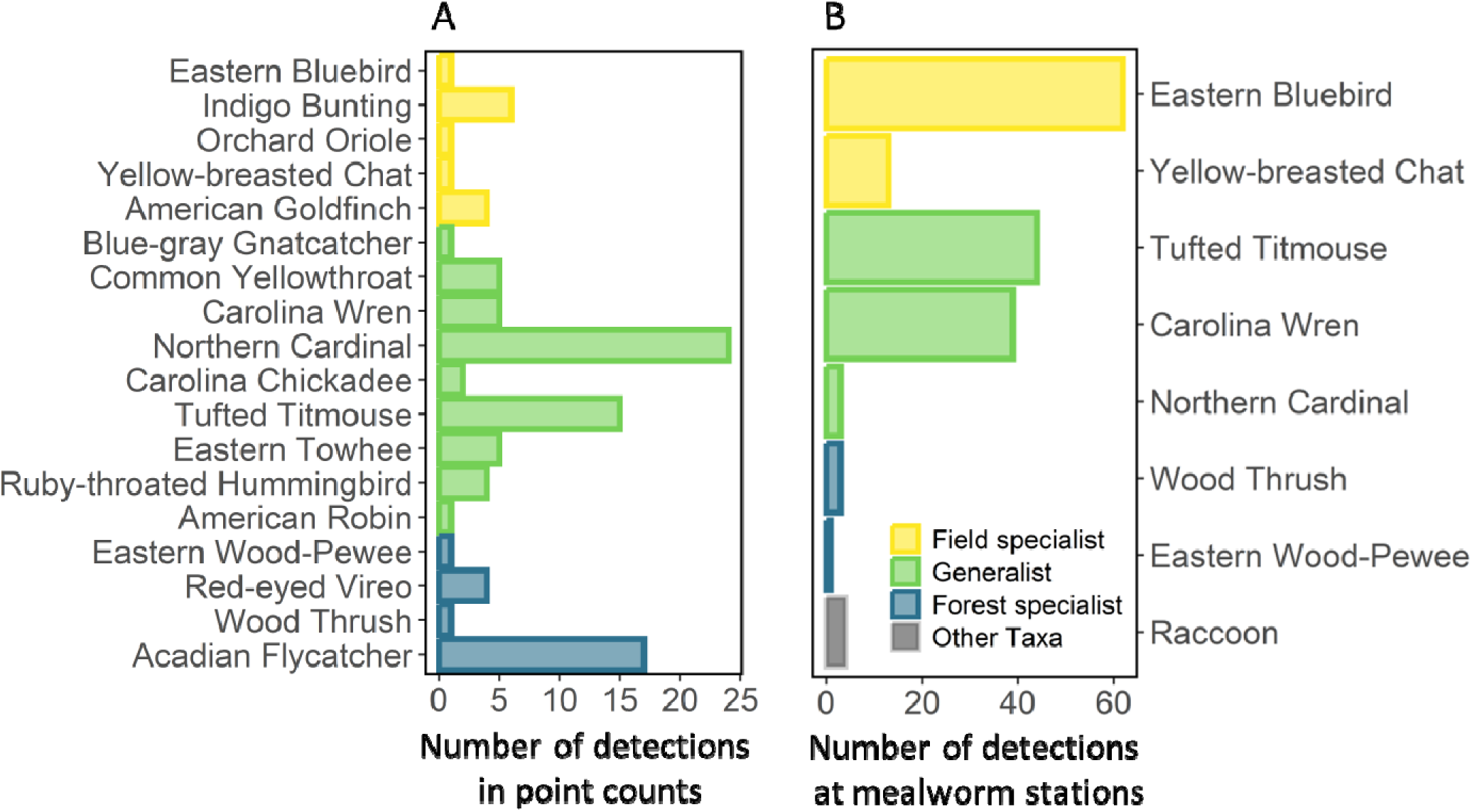
Although forest specialists and generalists were most commonly observed in plots, field specialists and generalists most frequently interacted with sentinel prey feeding stations. (A). Number of observations of each species observed in point counts in BiodiversiTREE plots. Colors indicate whether species were classified as field specialists (yellow), generalist (green), and forest specialists (blue-green) based on their relative abundances in NEON plots across the study landscape. (B) Number of camera trap detections (photos) of species visiting feeding stations with sentinel prey.

### Effect of forest structure, microclimate, and exclosures on insect herbivory

Global additive models of leaf herbivory showed that chewing, skeletonizing, and total insect herbivory all varied significantly across the three tree species sampled (BCIs for contrasts excluded 0; Appendix S1: Table 3). Herbivore damage was lowest on sweet gum (*L. styraciflua*) leaves, intermediate on beech (*F. grandifolia*), and greatest on white oak (*Q. alba*) leaves. The probability of leaf damage generally increased from the first survey (before placing exclosures) to the second survey (after exclosure removal) for both treatment and control branches (Appendix S1: Table 3). In additive models, there was no statistical support for the direct effect of tree diversity, canopy height, or bird detections on the probability of any of the leaf herbivory outcomes (BCIs of effect sizes included 0). However, there was statistical support for the effect of air temperature on the probability of total herbivory (β = -0.44; BCI = [-0.78, -0.04]) and skeletonizing herbivory (Fig. 4; β = -0.49, BCI = [-0.93, -0.002]), indicating that probability of leaf damage from herbivory decreased with increasing air temperature.

**Figure 4.**
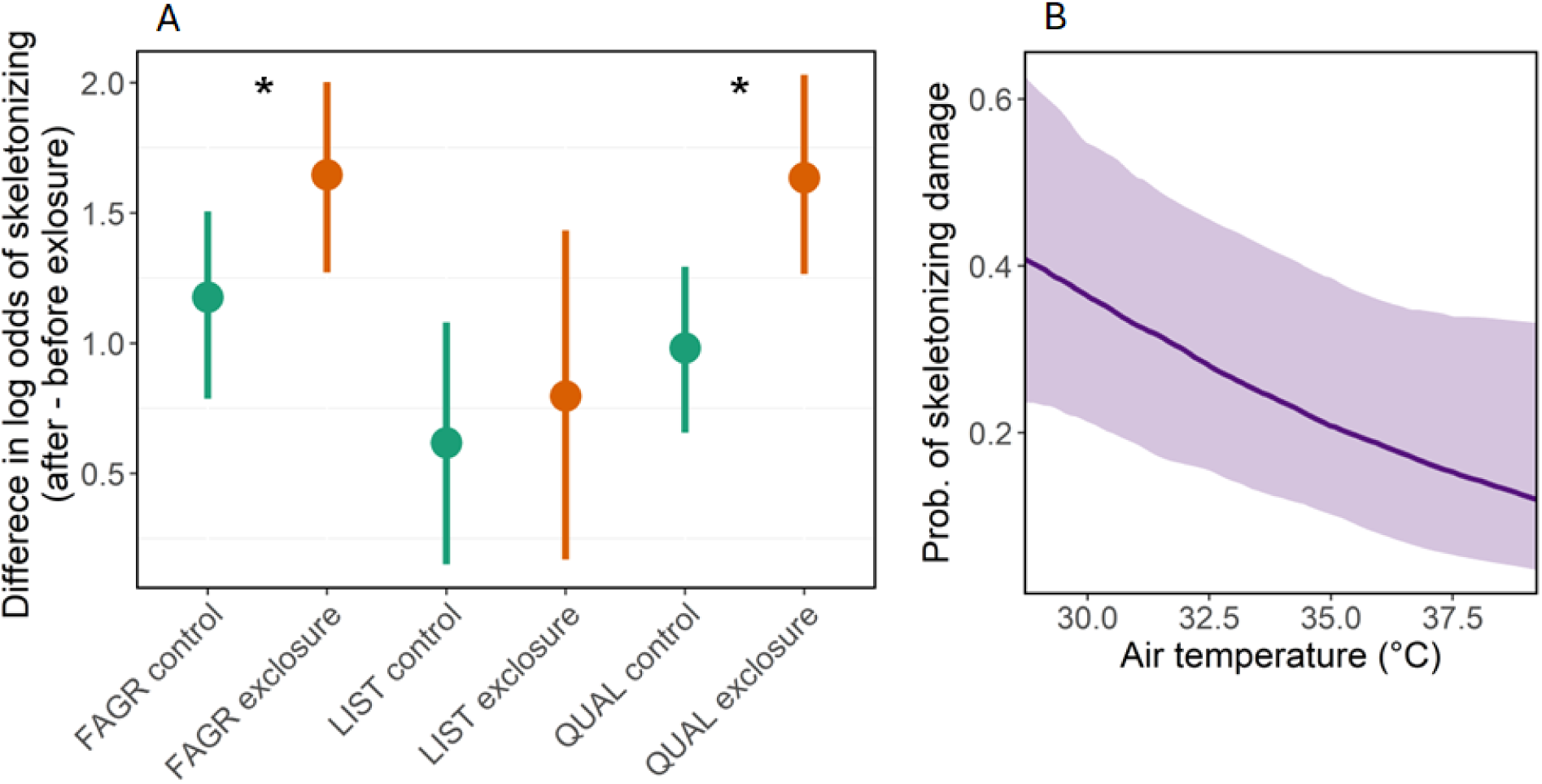
Skeletonizing patterns of leaf herbivory on American beech (*Fagus grandifolia*; FAGR), sweet gum (*Liquidambar styraciflua*; LIST) and white oak (*Quercus alba*; QUAL) increased following exclusion of bird predators and were more prevalent in cooler plots. (A) Points show difference in the log odds of skeletonizing patterns of leaf damage, before and after placement of exclosures. Greater values for exclosure branches than control branches indicate that this type of herbivory generally increased as a result of bird exclosures. Asterisks indicate significance of pairwise comparisons. Estimates were derived from the best supported interactive model in a set of Bayesian generalized linear models. Error bars are 95% Bayesian credible intervals. (B) Predicted effect of 95th percentile of near-ground daytime air temperatures on probability of skeletonizing patterns of leaf herbivory, based on a generalized linear model containing only additive terms. The bands indicate 95% credible intervals.

The most parsimonious interactive models (lowest LOOIC values; Appendix S1: Table 4) indicated statistical support for the effect of interactions between BACI effects (before/after, exclosure/control) and focal tree species on probability of leaf herbivory for total insect herbivory and skeletonizing herbivory response types. However, these interactions reflected tree species dependent increases in herbivory over the experiment (before/after) rather than exclosure effects (treatment/control). To isolate exclosure effects, accounting for overall changes in herbivory from the first to second leaf surveys (before/after), we first calculated the difference in posteriors for before and after estimates for each species and exclosure (treatment/control) combination. and then we calculated the difference-in-difference between enclosure and control estimates. These difference-in-difference calculations showed no support for the effect of exclosures on total herbivory or chewing herbivory (BCIs for contrasts include 0; Appendix S1: Fig. 4). However, there was support for the effect of exclosures on skeletonizing herbivory, where the probability of this type of herbivory significantly increased following the placement of bird exclosures, relative to the controls (Fig. 4; overall mean contrast across species = 1.30; BCI = [0.25, 2.23]). The interactive models did not indicate support for the dependence of treatment on environmental covariates (BCIs for estimates of interaction terms included 0).

## DISCUSSION

Numerous national and international initiatives promote tree planting as a nature-based solution aimed at climate change mitigation and maintaining local biodiversity (Di Sacco et al. 2021). Despite large monetary investments and increased research attention, we still lack a predictive framework for understanding how animal communities are reassembled in restoration areas, the interactions between restoration outcomes and climate change, and the consequences for ecosystem function. Recent research shows that the combination of microclimate exposure and climate niches of species can play a central role in restructuring assemblages across landscapes, including studies of ectotherm and bird assemblages (Frey, Hadley, and Betts 2016; Frishkoff et al. 2016; Nowakowski, Watling, et al. 2018; Nowakowski, Frishkoff, et al. 2018; Tourani et al. 2023). Here we find that microclimate is a significant driver of bird occurrence and leaf herbivory rates at fine spatial scales, across tens of meters. We also show that habitat and microclimate affiliation of birds modified predation risk to an insect herbivore, and top-down effects of bird exclosures were detected for skeletonizing patterns of leaf herbivory. The results suggest microclimate resulting from variation in tree diversity and forest structure can shape space use of birds at fine scales with consequences for community reassembly and trophic interactions in planted forests. These lessons can inform active forest restoration efforts that aim to recover biodiversity and associated ecosystem services.

### Fine-scale patterns of space use by birds

We addressed three interrelated questions to understand how fine-scale temperature variation and measures of forest structure influenced bird assemblages and associated trophic interactions within a forest restoration experiment. We first examined the effects of tree diversity, forest structure, and microclimate on fine-scale patterns of bird occurrence across plots. Both probability of occurrence and number of bird species detected increased in cooler plots. Neither tree diversity nor forest structure were significant predictors of bird occurrence after controlling for air temperature but were important for indirectly and directly mediating microclimate exposure (Fig 1). This result suggests that promoting cooler microclimates through management of tree diversity and structure may be important for improving outcomes for bird biodiversity in areas undergoing forest restoration.

High tree diversity within planted forest plots was associated with increased vertical forest structure based on LiDAR measurements, likely as a result of the variable growth rates and heights of planted tree species. In turn, greater maximum canopy height and vertical structure were both associated with cooler near-ground air temperatures within plots. A growing literature highlights the role of forest structure in buffering below-canopy temperatures and decoupling these microclimates from macroclimate conditions measured from weather stations in open areas—the primary data used in most ecological climate change assessments (Nowakowski, Frishkoff, et al. 2018; Zellweger et al. 2020). Research has demonstrated the direct influence of canopy height and openness, vertical structure, shade casting and composition of individual trees, and sub-canopy microhabitats on near-ground microclimate conditions (Chen et al. 1999; Jucker, Bouriaud, and Coomes 2015; Devaney et al. 2017; Lenoir, Hattab, and Pierre 2017; Nowakowski, Frishkoff, et al. 2018; Atkins et al. 2023; Gril et al. 2023; Verheyen et al. 2024).

Other studies demonstrate that canopy structure (height and openness) can depend on tree species diversity (Jucker, Bouriaud, and Coomes 2015; Ehbrecht et al. 2021). However, further research is needed to assess the relative importance of underlying measures of forest structure, the contribution of tree traits to cooling, and the responses of multiple dimensions of microclimate. Furthermore, most research to date has focused on microclimates within mature and secondary forests (Suggitt et al. 2011; von Arx et al. 2013; De Frenne et al. 2021; Kim et al. 2022), with less attention paid to microclimate changes under different restoration approaches and the consequences for the reassembly of forest communities following restoration.

After controlling for tree diversity and forest structure, microclimate was the most important direct driver of patterns of bird occurrence at fine spatial scales, across plots.

Temperature can be an important predictor of bird activity, such as breeding and foraging behaviors (Frey, Hadley, and Betts 2016; Kim et al. 2022), which are likely linked to species-specific thermal preferences and physiological thresholds. When exposed to low and high temperatures, beyond a species’ thermal neutral zone, individuals experience an increase in resting metabolic rate, with potential fitness costs (Khaliq et al. 2014). For example, greater energy expenditures when exposed to higher temperatures can reduce biomass of forest birds (Aggarwal et al. 2023), and exposure to extreme heat and drought can reduce survival (Iknayan and Beissinger 2018). These and other mechanisms associated with microclimate exposure can limit the distributions of individuals in space and time. Therefore, thermal trait-by-environment interactions likely filter species assemblages, as microclimates change under forest restoration.

Fine-scale space use by birds reflected landscape-scale patterns of occurrence. Specifically, the magnitude of the effect of microclimate on bird occurrences was partially dependent on landscape-scale habitat affiliations (field vs. forest) of species. Forest specialists at the landscape scale exhibited the strongest association with cooler microclimates at fine spatial scales. Whereas, habitat generalists and field specialists exhibited moderate and weak associations with microclimate, respectively (Fig 2). At global and continental scales, macroclimate gradients are a dominant factor shaping the distributions of biomes, communities, the geographic ranges of species, and patterns of diversity (Thomas 2010; Pyron and Wiens 2013; Ficetola et al. 2021). Interaction rates tend to increase in regions with greater species diversity, leading to increased predation and herbivory in the tropics (Lim, Fine, and Mittelbach 2015; S. Zhang, Zhang, and Ma 2016; Roslin et al. 2017; Freestone et al. 2021). Recent work has also shown that the macroclimate affiliations of species throughout their geographic ranges can predict their distributions across habitats within landscapes, indicating that climate affiliations scale from continents to landscapes (Frishkoff et al. 2016). Our results extend this scaling phenomenon by showing that fine-scale space use was also predicted by microclimate variation across tens of meters, with the strength of responses dependent on habitat affiliations across the landscape. Collectively, this evidence suggests that microclimate may be a fundamental determinant of species distributions, from local to landscape to global scales. By facilitating local persistence of species, enhancing microclimate refugia through restoration may ultimately slow large-scale distribution shifts under climate change (Zellweger et al. 2020; Huang et al. 2023).

### Predation risk across a microclimate gradient

We then asked whether predation risk for herbivorous insects by birds varied across gradients of tree diversity, forest structure, and microclimate. The results of the sentinel prey experiment showed that microclimate was also the most important environmental predictor of predation risk. However, despite greater probability of bird occurrence in cooler plots, bird predation of mealworms was greatest in warmer plots, running counter to our expectation that the increased occurrence of birds would also increase overall predation risk. Instead, we found that a subset of the bird assemblage—largely field-associated and generalist bird species—were disproportionately detected at mealworm stations (Fig 3) and were therefore likely responsible for most of the recorded predation.

Several factors may have contributed to increased predation in warmer plots by field-associated and generalist species. First, ectothermic insects are typically more active at higher temperatures due to elevated metabolism (Heinrich 1974; Dillon, Wang, and Huey 2010), and increased movement of mealworms in warmer plots may have attracted greater attention from birds. Second, differences in foraging behavior among common species in our surveys may have influenced observed spatial variation in predation risk. For example, some of the dominant forest specialist species observed in point counts were flycatchers that forage, in part, by hawking insects in flight or gleaning insects in the midstory or canopy (Allen et al. 2020; Watt et al. 2020), whereas some of the generalist and field specialist species detected in surveys, like Carolina Wrens and Eastern Bluebirds, spend more time foraging near the ground and were frequent visitors of feeding stations (Gowaty and Plissner 2020; Haggerty and Morton 2020). Third, many bird species are known to modify foraging behavior depending on tree species, vegetation structure, and prey availability (Holmes and Schultz 1988; Morrison et al. 2010; Aggarwal et al. 2023). For example, within forests subject to selective logging, birds spent more time foraging in warmer, logged plots where arthropod abundance was lower than in old growth forest (Aggarwal et al. 2023). The background prey availability also likely varied across plots during our experiment, as other studies in BiodiversiTree have found that spiders, caterpillars, and ground beetles were more abundant in plots with greater tree diversity, height, and canopy cover (Butz et al. 2023), Bennett, in prep)—i.e., those with cooler microclimates. We speculate that birds within warmer plots may have preferred mealworms to conserve energy that would otherwise be dedicated to foraging, given that mealworms were an accessible, added food source in these plots where natural prey insects were likely more limited than in cooler plots.

### Top down effects of bird predation on leaf herbivory

Last, we assessed whether top-down effects of avian predation of insect herbivores affected prevalence of leaf damage and whether these effects varied across gradients of tree diversity, forest structure, and microclimate. We found that the probability of all types of leaf damage varied significantly among focal tree species, with greatest leaf damage on white oaks. Variation in insect abundance across different tree species and levels of plant quality can impact bird-insect interactions; previous work has shown that greater insect abundances were associated with increased bird predation and more pronounced impacts of bird exclusion (Giffard et al. 2012; Singer et al. 2012). Other studies have found that arthropod abundance increases as tree species diversity increases, which is expected to drive subsequent increases in bird predation under the natural enemies hypothesis; yet, the evidence supporting this prediction is mixed (Giffard et al. 2012; Stemmelen et al. 2022). We did not find support for the natural enemies hypothesis in this study, as top-down effects of bird exclosures did not vary significantly across tree species or levels of tree diversity.

There is broad evidence that insectivorous birds often exert control over abundances of herbivorous insects in forests, reducing leaf damage that can in turn reduce the growth and survival of seedlings and mature plants (Mäntylä, Klemola, and Laaksonen 2011; Dekeukeleire et al. 2019; Getman-Pickering et al. 2023). Anthropogenic disturbances such as logging and fragmentation can modify the abundances of both predators and herbivores, potentially altering these interactions (Peter et al. 2015; Aggarwal et al. 2023). Here we found that total insect damage and skeletonizing herbivory decreased with increasing air temperature, with lowest herbivory occurring in open plots with less vertical structure. This result agrees with observations of greater abundances and diversity of some arthropod groups in cooler plots within the study site (Butz et al. 2023), Bennett et al. in prep). However, contrary to our expectations, we did not find that top-down effects of birds varied significantly across forest structure and microclimate gradients (i.e., no significant treatment-by-environment interaction). Although non-significant, there was a trend toward greater increases in herbivory on exclosure branches in cooler plots, as we would expect given greater bird diversity and herbivory in those plots. The length of the experiment could have limited our ability to detect drivers of spatial variation in top-down effects, which may also vary across seasons. Additionally, the exclosures could have prevented certain species of arthropods (e.g., larger Lepidopteran species) from ovipositing on treatment branches, potentially reducing total caterpillar abundance and associated patterns of herbivory. Furthermore, some bird species selectively forage for larger caterpillars (Singer et al. 2017) which could collectively contribute to differences in response to bird exclusion between chewing and skeletonizing patterns of herbivory.

## Conclusions

We found that microclimate was a dominant driver of fine-scale variation in bird occurrences, predation risk for an insect prey species, and leaf herbivory across planted forest plots. These findings add to a growing body of research that demonstrates the critical role of microclimate and the thermal ecology of organisms in shaping responses to global change drivers, including climate- and forest cover changes. Thermal trait-by-environment interactions can exert a strong influence on the distributions of birds across landscapes (Frey, Hadley, and Betts 2016) and at fine scales (this study), the abundances and composition of insects (Williamson et al. 2022, Bennett, in prep), and the interactions between these groups (Aggarwal et al. 2023), this study). The control of microclimate over assemblages at multiple trophic levels holds important implications for the reassembly of communities in planted and regenerating forests, their consequent interactions, and the resulting effects on forest structure and ecosystem process like biodiversity maintenance and carbon sequestration.

Multiple international frameworks promote forest restoration as a key nature-based solution to climate change and intervention for maintaining biodiversity (IPCC 2019, Kunming-Montreal Global Biodiversity Framework. 2022). In this study, higher tree diversity indirectly cooled below-canopy microclimates through increased vertical forest structure, thereby promoting local diversity of birds visiting plots. Promoting local animal diversity can in turn affect forest structure, with potential to increase carbon sequestration, through seed dispersal, pollination, herbivory, and mycorrhizal diversity (Schmitz et al. 2023, Bello et al. 2024). These results highlight the importance of considering tree diversity and microclimate management in the growing number of tree planting and restoration initiatives, globally. There are opportunities to optimize the effectiveness of forest restoration for biodiversity outcomes by planting mixtures of species that enhance cooling and integrating knowledge of microclimate connectivity and refugia into spatial planning. Closing this research-implementation gap will facilitate both the maintenance and adaptation of biodiversity under climate change.

## Supporting information

Supplemental Tables and Graphs

## Acknowledgments

SC was supported by an NSF Research Experiences for Undergraduates site grant (DBI-2244132). EO received support from a gift in honor of Alan Ullberg. This research was further supported by a Smithsonian Institution Scholarly Studies Award and a generous gift from John C. and Ann Ryan.

## Author Contributions

SC, EO, SKB, JDP, and AJN designed the study. SC, EO, and AJN analyzed the data. All authors collected data and contributed to edits of the manuscript.

