## Supplemental Tables and Graphs for "Bird occurrence and trophic interactions vary across gradients of tree diversity and microclimate in a planted forest"

### APPENDIX

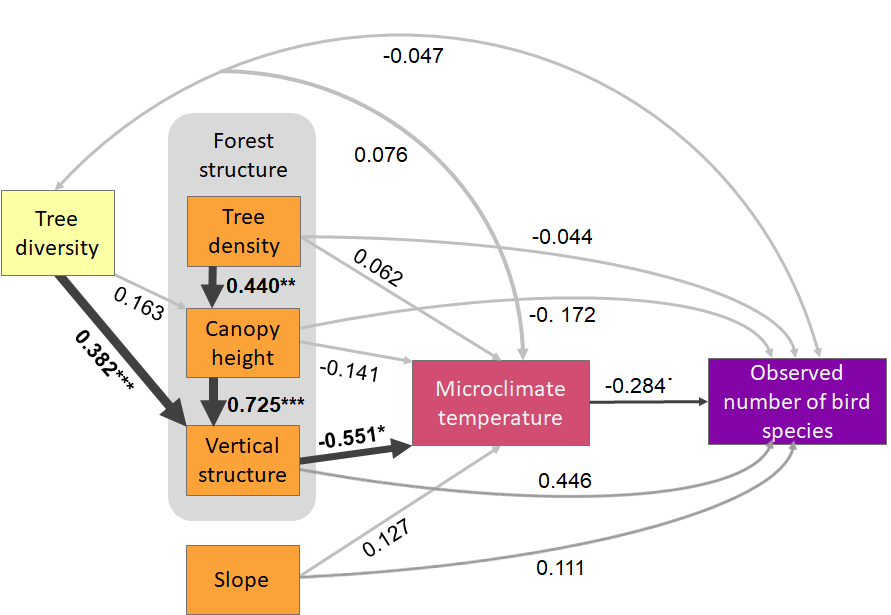

Fig. S1. After controlling for diversity, structure, and topography and although marginally non-significant, microclimate was a primary driver of variation in the number of observed bird species across plots. Diagram shows hypothesized causal pathways among tree diversity, forest structure, slope, microclimate and probability of bird occurrences in forest plots. Values shown are standardized coefficients from a piecewise structural equation model. Black arrows indicate significant model estimates for a given path ( ^.^*P* < 0.10, **P* < 0.05, ***P* < 0.01, ****P* < 0.001).

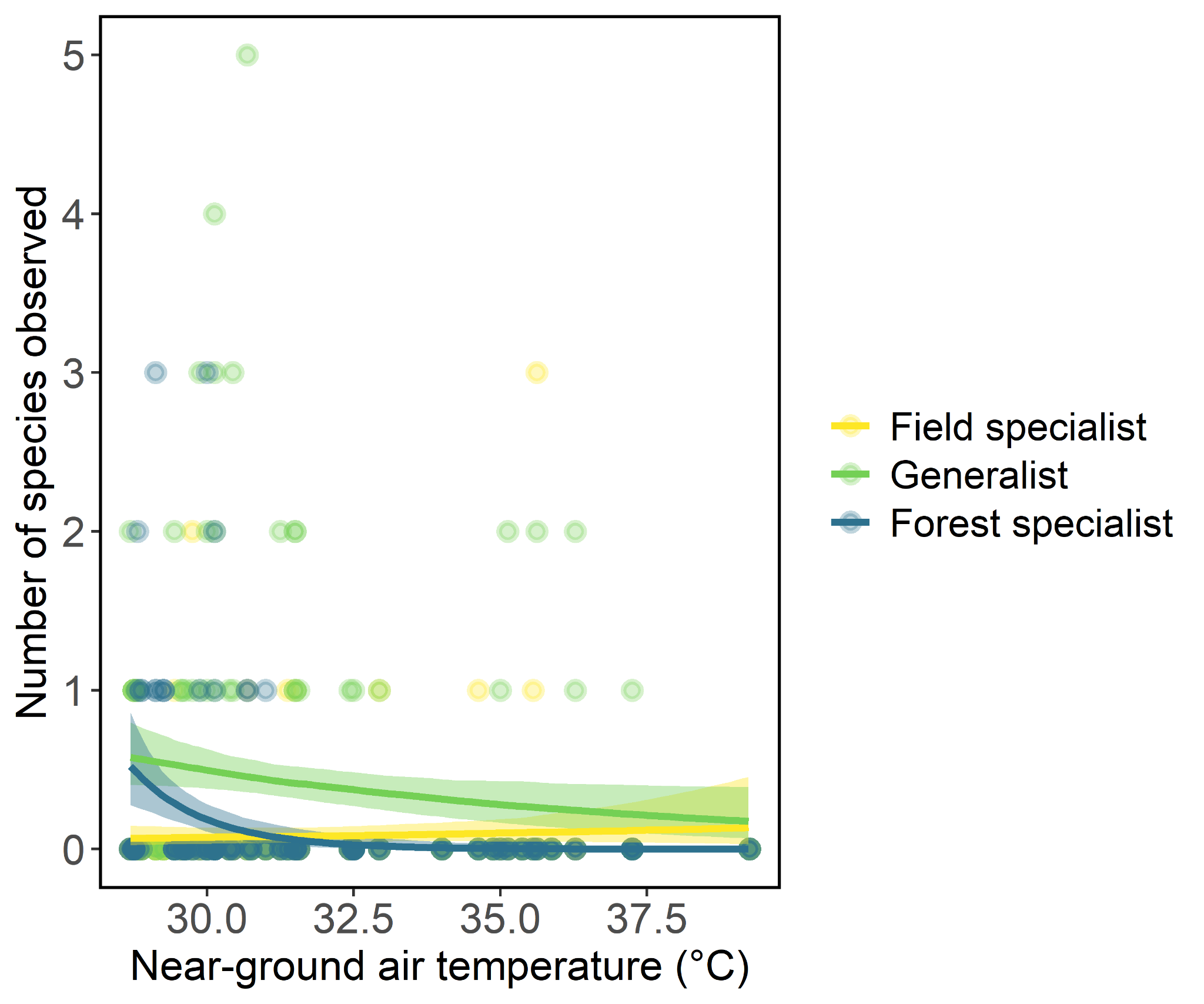

Fig. S2. Responses of birds to microclimate depended partially on their macrohabitat affiliations. The plot shows the predicted effect of the interaction between landscape habitat affiliations and responses to microclimate variation at fine spatial scales across plots, from a generalized linear model. Bands indicate 95% credible intervals for predictions.

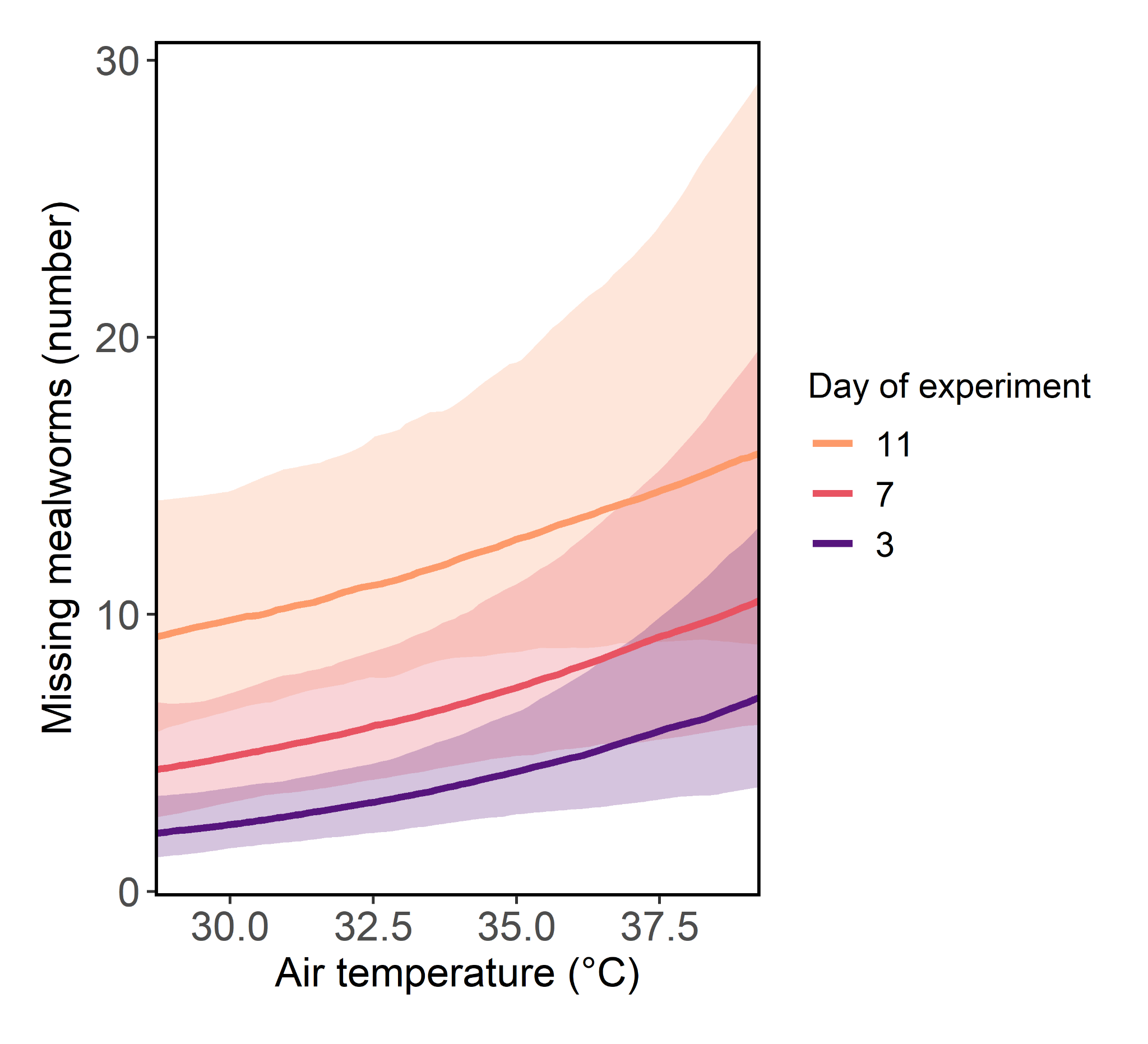

Fig. S3. Rates of mealworm loss increased with increasing plot temperature and over the duration of the experiment. The predicted interactive effect of 95th percentile of near-ground daytime air temperatures and day of experiment on the number of missing mealworms is based on a generalized linear mixed model. The bands indicate 95% credible intervals.

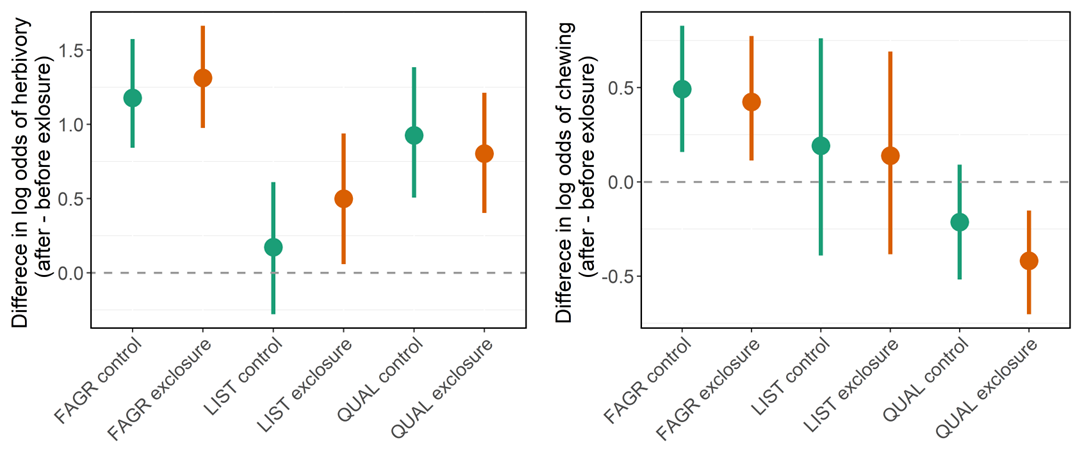

Fig. S4. Total insect herbivory (A) and chewing (B) patterns of leaf herbivory on American beech (*Fagus grandifolia*; FAGR), sweet gum (*Liquidambar styraciflua*; LIST) and white oak (*Quercus alba*; QUAL) did not differ significantly between control and exclosure branches. Points show difference in the log odds of leaf herbivory, before and after placement of exclosures. Asterisks indicate significance of pairwise comparisons. Estimates were derived from Bayesian generalized linear mixed models. Error bars are 95% Bayesian credible intervals.

Table S1. Responses of birds to microclimate depended partially on their macrohabitat affiliations. The table includes estimates from Bayesian Generalized Linear Models examining the response of bird occurrence or number of species to the interaction between landscape habitat affiliations and microclimate variation at fine spatial scales across plots. Bands indicate 95% credible intervals for predictions. The table includes the standard error (SE), lower (LCI) and upper (UCI) values of the 95% Credible Intervals, and the Gelman-Rubin statistic (Rhat) for each parameter.

| **Response** | **Model term** | **Estimate** | **SE** | **95% LCI** | **95% UCI** | **Rhat** | **Sig** |
| --- | --- | --- | --- | --- | --- | --- | --- |
| Occurrence | Intercept | -3.45 | 0.71 | -5.02 | -2.34 | 1.00 | * |
| Occurrence | Temperature | -2.53 | 0.81 | -4.47 | -1.24 | 1.00 | * |
| Occurrence | Habitat (Generalist) | 2.38 | 0.74 | 1.20 | 4.00 | 1.00 | * |
| Occurrence | Habitat (Field specialist) | 0.72 | 0.78 | -0.66 | 2.35 | 1.00 |  |
| Occurrence | Temperature:Habitat (Generalist) | 2.14 | 0.83 | 0.73 | 4.06 | 1.00 | * |
| Occurrence | Temperature:Habitat (Field specialist) | 2.57 | 0.89 | 1.16 | 4.58 | 1.00 | * |
| Species | Intercept | -3.10 | 0.55 | -4.37 | -2.16 | 1.00 | * |
| Species | Temperature | -2.14 | 0.62 | -3.50 | -1.08 | 1.00 | * |
| Species | Habitat (Generalist) | 2.18 | 0.56 | 1.20 | 3.45 | 1.00 | * |
| Species | Habitat (Field specialist) | 0.58 | 0.62 | -0.54 | 1.93 | 1.00 |  |
| Species | Temperature:Habitat (Generalist) | 1.82 | 0.63 | 0.70 | 3.22 | 1.00 | * |
| Species | Temperature:Habitat (Field specialist) | 2.32 | 0.67 | 1.15 | 3.72 | 1.00 | * |

Table S2. Rates of mealworm predation increased with both temperature and time. Estimates are from a Bayesiean implementation of a Generalized Linear Mixed Model with the number of mealworms missing from feeding stations as the responses variable. Plot was included as a varying intercept. The table includes the standard error (SE), lower (LCI) and upper (UCI) values of the 95% Credible Intervals, and the Gelman-Rubin statistic (Rhat) for each parameter.

| **Variable** | **Estimate** | **SE** | **95% LCI** | **95% UCI** | **Rhat** | **Sig** |
| --- | --- | --- | --- | --- | --- | --- |
| Intercept | 1.780 | 0.175 | 1.439 | 2.124 | 0.997 | * |
| Temperature | 0.265 | 0.107 | 0.042 | 0.473 | 0.998 | * |
| Day of experiment | 0.624 | 0.052 | 0.527 | 0.727 | 0.999 | * |
| Canopy height | 0.131 | 0.112 | -0.073 | 0.365 | 0.998 |  |
| Vertical structure | -0.026 | 0.138 | -0.286 | 0.252 | 0.998 |  |
| Tree diversity (4 spp) | 0.213 | 0.273 | -0.288 | 0.752 | 0.999 |  |
| Tree diversity (12 spp) | 0.188 | 0.248 | -0.301 | 0.692 | 1.005 |  |
| Temperature:Day | -0.094 | 0.032 | -0.157 | -0.029 | 1.001 | * |
| Canopy height:Day | -0.063 | 0.033 | -0.124 | 0.001 | 0.999 |  |
| Vertical structure:Day | 0.006 | 0.036 | -0.062 | 0.070 | 1.003 |  |
| Tree diversity (4 spp):Day | -0.071 | 0.071 | -0.215 | 0.068 | 1.005 |  |
| Tree diversity (12 spp):Day | -0.118 | 0.073 | -0.259 | 0.029 | 1.001 |  |

Table S3. Estimates from additive models of the effects of exclosures, tree species, tree diversity, and canopy height, temperature, and bird detections during point counts on leaf herbivory. The table includes the standard error (SE), lower (LCI) and upper (UCI) values of the 95% Credible Intervals, and the Gelman-Rubin statistic (Rhat) for each parameter.

| **Response** | **Model term** | **Estimate** | **SE** | **95% LCI** | **95% UCI** | **Rhat** | **Sig** |
| --- | --- | --- | --- | --- | --- | --- | --- |
| Herbivory | Intercept | -0.108 | 0.329 | -0.733 | 0.545 | 0.999 |  |
| Herbivory | After - Control | 0.789 | 0.107 | 0.584 | 0.995 | 1.003 | * |
| Herbivory | Before - Exclosure | -0.178 | 0.248 | -0.695 | 0.317 | 1.007 |  |
| Herbivory | After - Exclosure | 0.716 | 0.246 | 0.231 | 1.175 | 1.000 | * |
| Herbivory | Species (LIST) | -2.162 | 0.356 | -2.801 | -1.441 | 1.003 | * |
| Herbivory | Species (QUAL) | 1.611 | 0.350 | 0.933 | 2.322 | 1.001 | * |
| Herbivory | Tree diversity (4 spp) | 0.061 | 0.288 | -0.482 | 0.619 | 1.000 |  |
| Herbivory | Tree diversity (12 spp) | 0.026 | 0.303 | -0.523 | 0.597 | 1.010 |  |
| Herbivory | Canopy height | -0.215 | 0.173 | -0.556 | 0.097 | 0.998 |  |
| Herbivory | Temperature | -0.440 | 0.192 | -0.777 | -0.041 | 0.998 | * |
| Herbivory | Bird detections | -0.021 | 0.171 | -0.349 | 0.328 | 1.002 |  |
| Chewing | Intercept | -1.336 | 0.212 | -1.741 | -0.922 | 0.999 | * |
| Chewing | After - Control | 0.088 | 0.113 | -0.127 | 0.300 | 1.003 |  |
| Chewing | Before - Exclosure | 0.071 | 0.164 | -0.240 | 0.392 | 1.003 |  |
| Chewing | After - Exclosure | 0.053 | 0.172 | -0.251 | 0.401 | 1.008 |  |
| Chewing | Species (LIST) | -1.247 | 0.223 | -1.670 | -0.802 | 1.005 | * |
| Chewing | Species (QUAL) | 1.334 | 0.221 | 0.911 | 1.730 | 0.998 | * |
| Chewing | Tree diversity (4 spp) | -0.202 | 0.194 | -0.563 | 0.169 | 0.998 |  |
| Chewing | Tree diversity (12 spp) | 0.280 | 0.187 | -0.081 | 0.633 | 1.002 |  |
| Chewing | Canopy height | -0.150 | 0.111 | -0.374 | 0.063 | 1.000 |  |
| Chewing | Temperature | -0.051 | 0.115 | -0.284 | 0.153 | 0.998 |  |
| Chewing | Bird detections | -0.026 | 0.111 | -0.243 | 0.205 | 1.001 |  |
| Skeletonizing | Intercept | -0.933 | 0.382 | -1.683 | -0.166 | 1.002 | * |
| Skeletonizing | After - Control | 1.002 | 0.106 | 0.806 | 1.214 | 0.999 | * |
| Skeletonizing | Before - Exclosure | -0.514 | 0.300 | -1.057 | 0.036 | 1.004 |  |
| Skeletonizing | After - Exclosure | 0.921 | 0.290 | 0.355 | 1.478 | 1.004 | * |
| Skeletonizing | Species (LIST) | -2.141 | 0.433 | -2.940 | -1.276 | 1.000 | * |
| Skeletonizing | Species (QUAL) | 0.698 | 0.432 | -0.066 | 1.583 | 0.999 |  |
| Skeletonizing | Tree diversity (4 spp) | 0.153 | 0.372 | -0.533 | 0.900 | 0.997 |  |
| Skeletonizing | Tree diversity (12 spp) | -0.174 | 0.400 | -1.005 | 0.576 | 1.006 |  |
| Skeletonizing | Canopy height | -0.104 | 0.215 | -0.570 | 0.319 | 1.007 |  |
| Skeletonizing | Temperature | -0.488 | 0.232 | -0.935 | -0.002 | 0.998 | * |
| Skeletonizing | Bird detections | 0.134 | 0.196 | -0.237 | 0.509 | 1.001 |  |

Table S4. Leave-one-out information criterion (LOOIC) for models of leaf herbivory with interaction terms between treatment and environmental variables hypothesized to mediate trophic interactions. The most parsimonious interactive models (lowest LOOIC values) indicated statistical support for the effect of interactions between BACI effects (before/after, exclosure/control) and focal tree species on probability of leaf herbivory for total insect herbivory and skeletonizing herbivory response types.

| **Response** | **Model terms** | **LOOIC** |
| --- | --- | --- |
| Herbivory | BACI*Species | 4242.1 |
| Herbivory | BACI*Species + BACI*Bird detections | 4239.0 |
| Herbivory | BACI*Species + BACI*Temperature | 4235.7 |
| Herbivory | BACI*Species + BACI*Canopy height | 4236.1 |
| Herbivory | BACI*Species + BACI*Tree richness | 4248.7 |
| Herbivory | BACI + Species + Birds + Canopy + Temp + Richness | 4252.5 |
| Chewing | BACI*Species | 4284.9 |
| Chewing | BACI*Species + BACI*Bird detections | 4285.7 |
| Chewing | BACI*Species + BACI*Temperature | 4290.8 |
| Chewing | BACI*Species + BACI*Canopy height | 4290.9 |
| Chewing | BACI*Species + BACI*Tree richness | 4289.2 |
| Chewing | BACI + Species + Birds + Canopy + Temp + Richness | 4301.5 |
| Skeletonizing | BACI*Species | 4023.1 |
| Skeletonizing | BACI*Species + BACI*Bird detections | 4018.5 |
| Skeletonizing | BACI*Species + BACI*Temperature | 4022.8 |
| Skeletonizing | BACI*Species + BACI*Canopy height | 4023.3 |
| Skeletonizing | BACI*Species + BACI*Tree richness | 4024.9 |
| Skeletonizing | BACI + Species + Birds + Canopy + Temp + Richness | 4022.6 |
